# A motion aftereffect from viewing other people’s gaze

**DOI:** 10.1101/2020.11.08.373308

**Authors:** William Randall, Arvid Guterstam

## Abstract

Recent work suggests that our brains may generate subtle, false motion signals streaming from other people to the objects of their attention, aiding social cognition. For instance, brief exposure to static images depicting other people gazing at objects made subjects slower at detecting subsequent motion in the direction of gaze, suggesting that looking at someone else’s gaze caused a directional motion adaptation. Here we confirm, using a more stringent method, that viewing static images of another person gazing in a particular direction, at an object, produced motion aftereffects in the opposite direction. The aftereffect was manifested as a change in perceptual decision threshold for detecting left versus right motion. The effect disappeared when the person was looking away from the object. These findings suggest that the attentive gaze of others is encoded as an implied agent-to-object motion that is sufficiently robust to cause genuine motion aftereffects, though subtle enough to remain subthreshold.

## Introduction

The ability of social animals to effortlessly track other’s attentional state is vital for constructing a theory of mind and predicting behavior (Baron-Cohen, 1997; Calder et al., 2002; Graziano and Kastner, 2011; Kobayashi and Kohshima, 1997). Recent evidence suggests that people construct a rich, implicit model of other people’s attention that goes far beyond merely registering the direction of gaze (Guterstam and Graziano, 2020a, 2020b; Guterstam et al., 2019, 2020a, 2020b; Kelly et al., 2014; Pesquita et al., 2016). This implicit model draws on the useful but physically highly inaccurate construct of beams of motion that emanate from the eyes and travel through space toward the attended object. This model is subthreshold. People generally show no explicit awareness of it, even though it has a significant effect on motion processing areas of the brain (Guterstam et al., 2020a), on behavioral measures of motion processing (Guterstam and Graziano, 2020a; Guterstam et al., 2019), and on social cognitive decisions about the attention of others (Guterstam and Graziano, 2020b). In this study, we made use of a visual phenomenon called the motion aftereffect to test a prediction of this proposed model: viewing static images depicting other people gazing in a particular direction, at an object, should lead to an illusory subsequent motion in the opposite direction.

The motion aftereffect is a classic phenomenon of a false motion signal in the visual image caused by prior exposure to motion in the opposite direction (Anstis et al., 1998; Wohlgemuth, 1911). It is typically assessed experimentally by first exposing subjects to a motion stimulus, including implied motion (e.g., a static image of a running animal) (Kourtzi and Kanwisher, 2000; Krekelberg et al., 2003; Winawer et al., 2008), and then measuring subjects’ speed and accuracy at detecting subsequent random-dot motion test probes (Glasser et al., 2011; Levinson and Sekuler, 1974). A genuine motion aftereffect is associated with slower reaction times and decreased accuracy for motion test probes of the same directionality as the adapting stimulus, reflecting direction-specific neuronal fatigue affecting motion processing time and perceptual decision-making. In a series of seven behavioral experiments (Guterstam and Graziano, 2020a), we previously showed that brief exposure to static images depicting a person gazing in a particular direction, at an object, made subjects significantly slower at detecting subsequent motion in the direction of gaze, which is compatible with a motion aftereffect caused by gaze encoded as implied motion. The effect disappeared when the depicted person was blindfolded or looked away from the object, and control experiments excluded differences in eye movements or asymmetric allocation of covert attention as possible drivers of the effect. However, because the paradigm in (Guterstam and Graziano, 2020a) was primarily designed for analysis of reaction time rather than accuracy, the task was made easy and accuracy was close to ceiling (mean accuracy across experiments = 91%). Thus, that experiment showed only reaction time effects, and failed to reveal any meaningful accuracy effects. The goal of the present study was to examine if seeing someone else’s gaze direction caused enough of a motion aftereffect to shift subjects’ perceptual decisions about subsequent motion. The present experiment is therefore a conceptual replication of the previous studies, but using a different measure of the motion aftereffect to test whether the discovery is reliable and robust across methods.

To achieve this goal, we modified the motion adaptation paradigm described in (Guterstam and Graziano, 2020a), which was based on a random-dot motion direction discrimination task, to maximize the likelihood of detecting meaningful differences in accuracy. Subjects were tested using an online, remote platform (Prolific) (Palan and Schitter, 2018) due to restrictions on research imposed by the coronavirus epidemic. (See Materials and Methods for details of sample sizes and exclusion criteria). Just as in (Guterstam and Graziano, 2020a), in each trial, subjects were first exposed to an image depicting a face on one side of the screen, gazing at a neutral object, a tree, on the other side (Fig 1A). After 1.5 s, the face-and-tree image disappeared, and subjects saw a random dot motion stimulus in the space interposed between where the head and the tree had been. The stimulus was shown for 1.0 s. The proportion of dots moving coherently in one direction (dot coherence) varied across seven different levels. The coherence ranged from 30% of the dots moving left (and 70% moving randomly) to 30% moving right in increments of 10% (thus, the middle condition of 0% coherence had 100% of the dots moving randomly). After the dots disappeared, subjects made a forced-choice left-or-right judgement of the global direction of the moving-dots stimulus.

**Figure 1.**
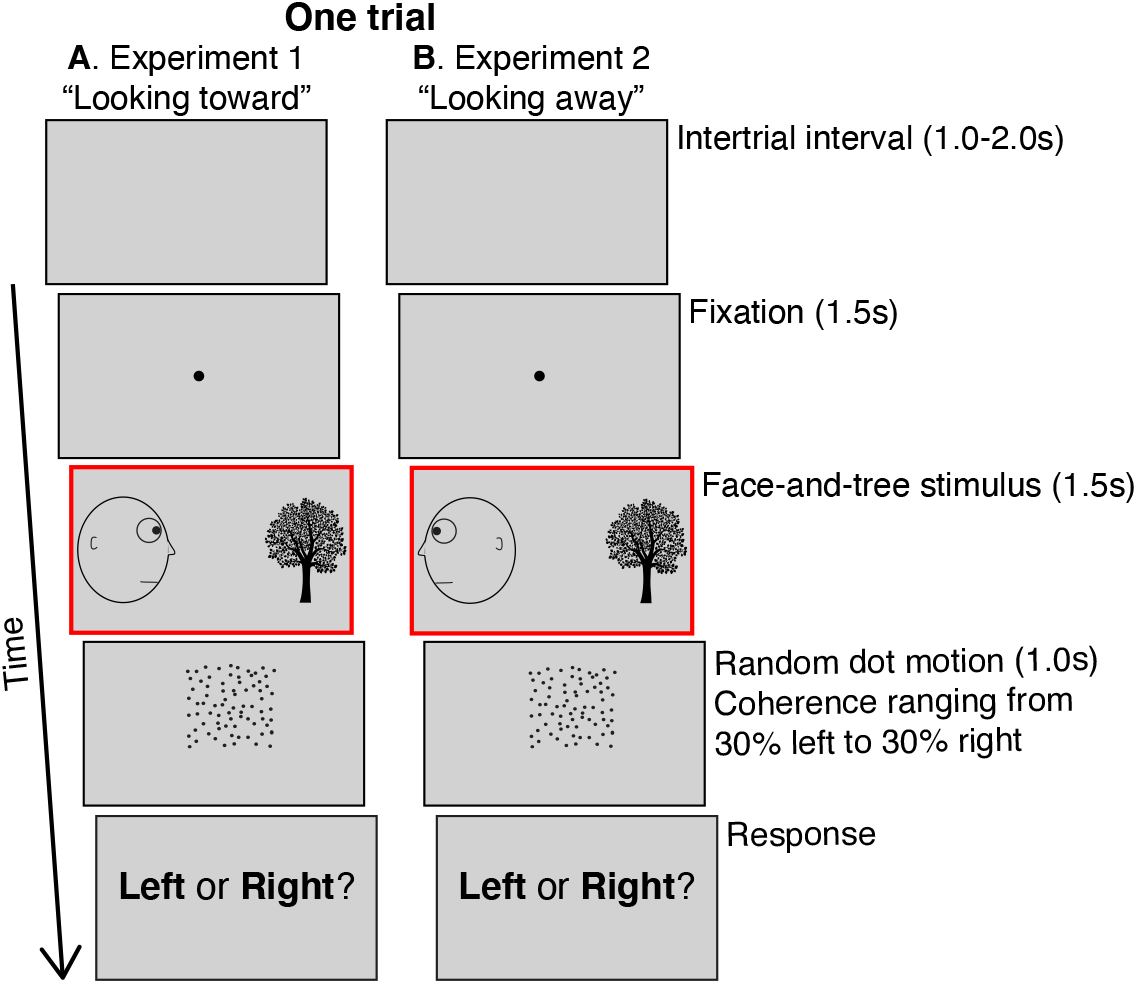
Behavioral paradigm. **A**. Experiment 1. After a 1-2 s inter-trial interval, subjects fixated on a central spot for 1.5 s, then saw a static, line-drawing image of a head looking at a tree for 1.5 s. The head could be on the left looking right (shown here), or on the right looking left. Subjects then saw a random dot motion stimulus for 1.0 s in the space interposed between where the head and the tree had been. A subset of the dots moved coherently either left or right. The dot coherence ranged from 30% of the dots moving left (and 70% moving in random directions) to 30% moving right, in 10% increments, yielding seven different types of motion. After the dots disappeared, subjects made a forced-choice judgement of the global direction of the moving-dots stimulus. **B**. Experiment 2. Similar to experiment 1 except the head was always looking away from the tree. The head could be on the left looking left (shown here), or on the right looking right. The red outline (not visible to subjects) indicate the key phase of the paradigm altered in experiment 1 versus 2.

This approach allowed us to calculate, at each level of coherence and on a subject-by-subject basis, the frequency of responses that were spatially congruent with the gaze direction in the preceding face-and-tree image (i.e., the direction toward the location of the tree). By fitting this data to a sigmoid function and extracting the sigmoid central point, we estimated the perceived null motion, that is, the amount of motion coherence for which subjects were equally likely to respond that the motion direction of the test probe was congruent or incongruent with the preceding gaze direction. We found that viewing another’s gaze significantly shifted the perceived null motion, as if that gaze caused an illusory motion aftereffect in the opposite direction (experiment 1). The effect disappeared when the face in the display was looking away from the object (experiment 2; Fig 1B), suggesting that the perception of the other actively gazing at the object was the key factor. These findings extend previous results by demonstrating that viewing other people’s gaze is associated with a false motion signal, below the level of explicit detection but still capable of generating a motion aftereffect that influences not only perceptual processing time, but also perceptual decision thresholds about subsequent motion.

## Results

In both experiment 1 (face looking toward the tree) and experiment 2 (face looking away from the tree), the appearance of the face on the left and tree on the right, or the face on the right and tree on the left, were balanced and presented in a random order. The subsequent dot-stimulus could move either leftward or rightward with 10%, 20% or 30% coherence, or be completely random (0% coherence). For analysis, the trial types were collapsed into seven conditions: -30%, -20%, -10%, 0%, +10%, +20%, and +30%, where motion toward the location of the (preceding) tree were arbitrarily coded as positive coherence, and motion away from the tree as negative coherence. Thus, the predicted motion aftereffect from viewing the face actively gazing in the direction toward the tree (in experiment 1) should produce a positive shift (>0%) of the perceived null motion. Subjects performed 70 trials in seven blocks of 10 trials each, thus 10 trials per condition.

Fig 2A and 2B summarize the results. In experiment 1 (n=59), where the face was looking at the tree, the central point of the sigmoidal function best describing subjects’ data across the seven different dot motion coherence conditions, was significantly greater than 0 (M = 1.18%, S.E.M. = 0.39%; t_58_ = 3.02, p = 0.0038; Fig 2B). This result confirmed our prediction that implied motion streaming from the eyes toward the tree causes a motion aftereffect in the opposite direction (Guterstam and Graziano, 2020a). In other words, immediately after subjects saw a face gazing one direction, the amount of real motion needed to make subjects think a test stimulus was randomly balanced between left and right movement was 1.18% coherence in the direction that the face had been gazing.

**Figure 2.**
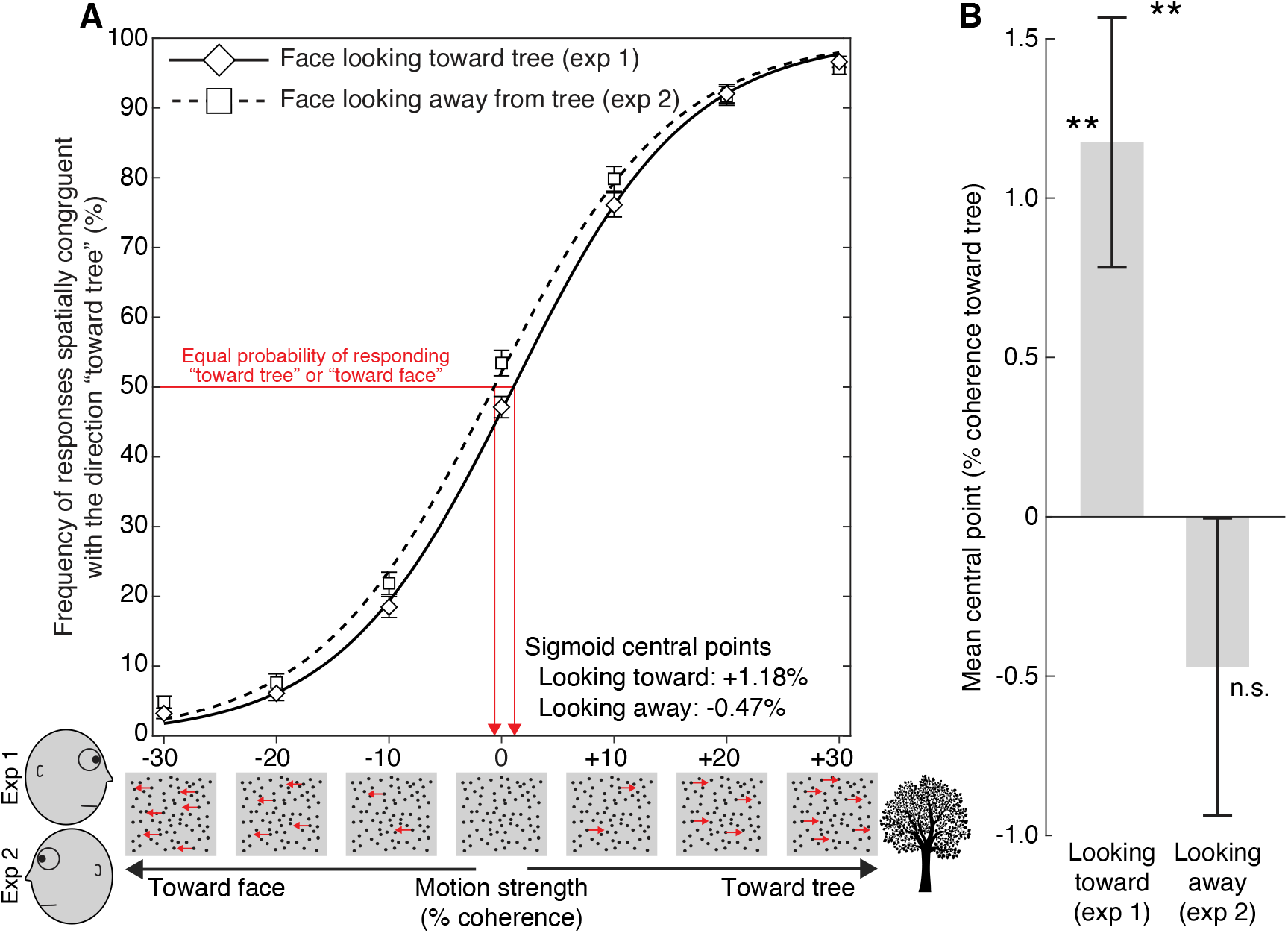
Results. **A**. For experiment 1 (“Face looking toward tree”; solid line) and experiment 2 (“Face looking away from tree”; dotted line), the frequency of dot-motion responses that were spatially congruent with the direction toward the tree is plotted as a function of motion coherence relative to the location of the tree. (We collapsed trials in which the tree appeared on the left or right side, and motion toward the tree is arbitrarily coded as positive coherence, and motion away from the tree as negative coherence.) Error bars indicate SEM. The sigmoid functions shown here were fitted to the group-mean values across coherence levels for the two experiments, for display purposes. **B**. Mean sigmoid central point estimates in experiment 1 and 2, based on subject-level fitting. The central point reflects the perceived null motion, that is, the amount of motion coherence for which subjects were equally likely to respond that the motion is “going toward the tree” as “going away from the tree”. When the face was looking at the tree (experiment 1), the central point was significantly greater than 0, and significantly greater compared to when the face was looking away from the tree (experiment 2). This pattern of results is compatible with the existence of a gaze-induced motion aftereffect in the space between agent and object. *p<0.05, **p<0.01. Error bars indicate SEM.

In experiment 2 (n=64), where the face was looking away from the tree, the central point of the sigmoid function was not significantly different from 0 (M = -0.47%, S.E.M. = 0.47%; t_63_ = -1.01, p = 0.3165; Fig 2B). In a between-groups comparison, we found that the central point was significantly modulated by the gaze direction of the preceding face (Looking toward [1.18%] versus Looking away [-0.47%]: t_121_ = 2.69, p = 0.0083; Fig 2B).

After all trials were completed, subjects in experiments 1 and 2 were asked what they thought the purpose of the experiment might be, and whether they were explicitly aware of any influence of the head-and-tree stimulus on their ability to respond to the dot motion stimulus. Though subjects offered guesses about the purpose of the experiment, none indicated anything close to a correct understanding. All subjects also insisted that, as far as they were aware, the head-and-tree stimulus had no impact on their response to the second stimulus. These questionnaire results suggest that any motion aftereffects observed here probably occurred at an implicit level.

## Discussion

These results strongly support the notion that when people view a face looking at an object, the brain treats that gaze as though a movement were present, passing from the face to the object. The motion test probes were more likely to be judged as moving in the direction opposite the gaze direction depicted in the previous adapting image than to be moving in the same direction, but only when the agent in the image was actively gazing at the object. This work extends previous results that focused on reaction times (Guterstam and Graziano, 2020a). Here, perception of other people’s gaze significantly biased perceptual decisions about subsequent motion, which is a hallmark of the motion aftereffect. We propose that this hidden motion signal, associated with gaze, is part of an implicit ‘fluid-flow’ model of other people’s attention, that assists in human social cognition.

The null result of experiment 2 suggest that spatial priming, i.e., subjects simply being more prone to choose the direction that the face was looking, is an unlikely explanation to the findings of experiment 1. Had spatial priming been the driving factor, one would expect a significant negative shift of the central point when the face was looking away from the object (experiment 2), of the same magnitude as the observed positive shift in experiment 1 where the face was gazing at the object. The absence of a motion aftereffect in experiment 2 also suggests that the presence of a face on one side of the screen drawing subjects’ attention or gaze more to that side, cannot easily explain effect observed in experiment 1.

The present set of results adds an important piece of evidence to a growing body of research on how people model the attention of others to support social cognition. The brain seems to represent other people’s attention as an implied, beam-like motion travelling from an agent to the attended object. This motion signal may be detected using sensitive behavioral motion adaptation paradigms, such as in the present study or in (Guterstam and Graziano, 2020a). It can also be quantified using a tube-tilting task, in which subjects’ angular judgements of the tipping point of a paper tube were implicitly biased by the presence of the gazing face, as if beams of force-carrying energy emanated from eyes, gently pushing on the paper tube (Guterstam et al., 2019). The motion signal is also detectable in brain activity patterns in the human motion-sensitive MT complex and in the temporo-parietal junction, which responded to the gaze of others, and to visual flow, in a similar manner (Guterstam et al., 2020a). Finally, by contaminating a subject’s visual world with a subthreshold motion that streams from another person toward an object, we could manipulate the subject’s perception of that other person’s attention, suggesting that subthreshold motion plays a functional role in social cognition (Guterstam and Graziano, 2020b).

Together, these present and previous findings suggest that the visual motion system is used to facilitate social brain mechanism for tracking the attention of others. We speculate that this implicit social-cognitive model, borrowing low-level perceptual mechanisms that evolved to process physical events in the real world, may help to explain the extraordinary cultural persistence of the belief in extramission, the myth that vision is caused by something beaming out of the eyes (Gross, 1999; Piaget, 1979; Winer et al., 1996).

## Materials and Methods

### Participants

For each experiment, participants were recruited through the online behavioral testing platform Prolific (Palan and Schitter, 2018). Using the tools available on the Prolific platform, we restricted participation such that no subject could take part in more than one experiment. Thus, all subjects were naÏve to the paradigm when tested. All participants indicated normal or corrected-to-normal vision, English as a first language, and no history of mental illness or cognitive impairment. All experimental methods and procedures were approved by the Princeton University Institutional Review Board, and all participants confirmed that they had read and understood a consent form outlining their risks, benefits, compensation, and confidentiality, and that they agreed to participate in the experiment. Each subject completed a single experiment in a 6-8 min session in exchange for monetary compensation. As is standard for online experiments, because of greater expected variation than for in-lab experiments, relatively large numbers of subjects were tested. A target sample size of 100 subjects per experiment was chosen arbitrarily before data collection began. Because of stringent criteria for eliminating those who did not follow all instructions or showed poor task performance (see below), initial, total sample sizes were larger than 100 and final sample sizes for those included in the analysis varied between experiments (experiment 1, n_total_=115, n_included_=59, 17 females, mean age 26 [SD 8]; experiment 2, n_total_=109, n_included_=64, 24 females, mean age 31 [SD 11]).

### Exclusion criteria

Subjects were excluded based on three predefined criteria: (i) poor task performance, (ii) poor curve fit, or (iii) failure read the instructions carefully. On average across the two experiments, the exclusion rate was 45% (experiment 1, n_excluded_=56 [49%]; experiment 2, 45 [41%]). The most common reason for exclusion was poor performance on the dot motion task (experiment 1, 44 [38%]; experiment 2, 34 [31%]). Because the task is meaningless if a subject cannot detect motion direction even at the easiest (highest) coherence levels, we excluded all subjects whose accuracy was less than 80% when 30% of the dots moved either right or left, in accordance with the exclusion criterium used in (Guterstam and Graziano, 2020a). The relatively high rate of exclusion due to poor performance here (35% on average) was expected given that the average exclusion rate was 19% in a previous study (Guterstam and Graziano, 2020a) using the same dot motion direction discrimination task but with a fixed 40% coherence level, which is easier to detect. Moreover, participants in (Guterstam and Graziano, 2020a) underwent up to four sets of 10 practice trials, with feedback, before commencing the main experiment, since reaction times (RTs), and not accuracy, was the outcome of interest in that study. In the present study, subjects did not undergo any practice sessions, because accuracy was our primary outcome. It therefore seems probable that the absence of practice trials and lower dot coherence levels in the present study fully explain the higher exclusion rates reported here compared to in (Guterstam and Graziano, 2020a).

Because the goal of the study was to estimate each individual’s sigmoid central point, which is disproportionally affected by a poor curve fit, we excluded subjects showing a goodness of fit (R^2^) below 0.9 (experiment 1, 12 [10%]; experiment 2, 11 [10%]). The mean goodness of fit among the included subjects was 0.97 in both experiment 1 and in experiment 2. The mean goodness of fit among those excluded were 0.80 in experiment 1 and 0.84 in experiment 2.

No subjects were excluded for failure to carefully read the instructions, which was determined by an instructional manipulation check (IMC). The IMC, also used in (Guterstam and Graziano, 2020b), was adapted from (Oppenheimer et al., 2009) and consisted of the following sentence inserted at the end of the instructions page: *“In order to demonstrate that you have read these instructions carefully, please ignore the ‘Continue’ button below, and click on the ‘x’ to start the practice session*.*”* Two buttons were presented at the bottom of the screen, “Continue” and “x”, and clicking on “Continue” resulted in a failed IMC.

### Apparatus

After agreeing to participate, subjects were redirected to a website where stimulus presentation and data collection were controlled by custom software based on HTML, CSS, JavaScript (using the jsPsych javascript library (de Leeuw, 2015)), and PHP. Subjects were required to complete the experiment in full screen mode. Exiting full screen resulted in the termination of the experiment and no payment. Because the visual stimuli were rendered on participants’ own web browsers, viewing distance, screen size, and display resolutions varied. The face-and-tree image encompassed 60% of the subject’s total screen width. Below, we report the stimulus dimensions using pixel [px] values for a screen with a horizontal resolution of 1050 px.

### Experiment 1

The experiment consisted of an initial instructions page, followed by the experimental session, and a post-experiment survey. The written instructions were as follows: “*Humans have an extraordinary ability to detect subtle visual motion. In this experiment, your task is to determine the motion direction of a field of small dots. Each trial consists of four phases, as indicated by the image below [subjects were presented an image featuring four sample frames indicating the four phases: (1) Fixation - “Keep looking in this spot throughout the trial”, (2) Image, (3) Dot motion, and (4) Response]. After viewing the dots for 1 sec, indicate the direction of the motion using the left or right arrow keys. There is always one correct answer. However, the motion direction will be obvious in some trials, and very subtle in other trials. The preceding face-and-tree image is irrelevant to the task and does not predict the motion direction in any way*.

Fig 1A shows the behavioral paradigm for experiment 1. After a variable 1-2 s inter-trial interval in which a neutral gray (R: 210, G: 210, B:210) field covered the screen, a black central fixation point (25px diameter) appeared. Subjects were instructed to fixate on the point and try to maintain fixation in that area of the screen throughout the trial. After 1.5 s, the point disappeared and a static image of a face (228 px wide and 250 px high) gazing across the screen toward an arbitrary object, a tree (196 px wide and 250 px high), was presented for 1.5 s.

When this static image disappeared, a random dot motion stimulus was presented for 1.0 s in the space interposed between where the head and the tree had been, vertically centered relative the position of the eyes of the face. The random dot motion stimulus featured black dots. Dot diameter was 2px. There were 400 dots within a 230 × 230 px square. Dot speed was 2 px/frame. Dot lifetime was 12 frames, which corresponds to 200ms on a standard 60 Hz monitor. There were seven different types of dot motion coherence levels: either 30%, 20%, or 10% of the dots moved leftward (while the 70%, 80%, or 90% of the dots moved in random directions), or 30%, 20%, or 10% of the dots moved rightward. In one condition, zero % of the dots moved coherently (i.e., 100% random directions). After 1.0 s, the dots disappeared, and the subjects were prompted about the direction of the motion (“Left or Right?). They indicated their response by pressing the left or right arrow key on their keyboard, after which the trial terminated and the next trial began.

For the statistical analysis, the trial types were collapsed into seven conditions: -30%, -20%, -10%, 0%, +10%, +20%, and +30%. Motion toward the location of the (preceding) tree were arbitrarily coded as positive coherence, and motion away from the tree as negative coherence. On a subject-per-subject basis, for each condition, we calculated the proportion of responses that was spatially congruent with the direction away from the face and toward the tree (which, in experiment 1, corresponded to the gaze direction of the face). We then fit the accuracy data to a sigmoidal function (Eq. 1) (Noel et al., 2020) using the Curve Fitting Toolbox for MATLAB (MathWorks). The unbiased value of 0.5 was used as starting points for the estimations of *x*_c_ and b in Eq. 1. For each subject, we extracted the central point of the sigmoidal (*x*_c_ in Eq. 1), representing level of dot coherence at which a subject is equally probable to indicate that the motion was moving toward the tree as compared to away from the tree.

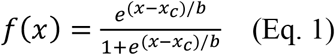

If the visual input of another’s active gaze has a motion aftereffect in the opposite direction, as predicted in experiment 1, then the central point should be significantly greater than 0. At the group level, we therefore compared the mean central point to 0 using a one-sample two-tailed t-test.

After the experiment, the subjects completed a survey. They were first asked an open-ended question: *“What do you think the hypothesis of the experiment was?”*. They were then given the binary yes-or-no question: “*Did the head-and-tree image influence your responses to the dots?”* Subjects who responded “yes” were asked, *“Please describe in what way the head-and-tree image influenced your responses to the dots*.*”* Subjects who in any way indicated that they had figured out the purpose of the experiment were excluded from the analysis.

### Experiment 2

The design, procedures, and statistical analysis of experiment 2 were identical to those of experiment 1, with one exception: the face was turned away from the tree. This control condition should eliminate any gaze-induced effect on motion judgments (Guterstam and Graziano, 2020a). We therefore predicted that the mean central point in experiment 2 would not significantly differ from to 0 (two-tailed one-sample t-test), and that it would be significantly smaller than the mean central point among participants in experiment 1 (two-tailed two-sample t-test).

## Acknowledgments

We thank Michael S. A. Graziano for valuable comments on an earlier version of the manuscript. This work was supported by the Princeton Neuroscience Institute Innovation Fund. Arvid Guterstam was supported by the Wenner-Gren Foundation, the Sweden-America Foundation, and the Promobilia Foundation.

## Data and materials availability

The data that support the findings of this study are available at https://figshare.com/s/f255024b75f884d7ec6f.

